## Supplementary material for "Intracerebral mechanisms explaining the impact of incidental feedback on mood state and risky choice"

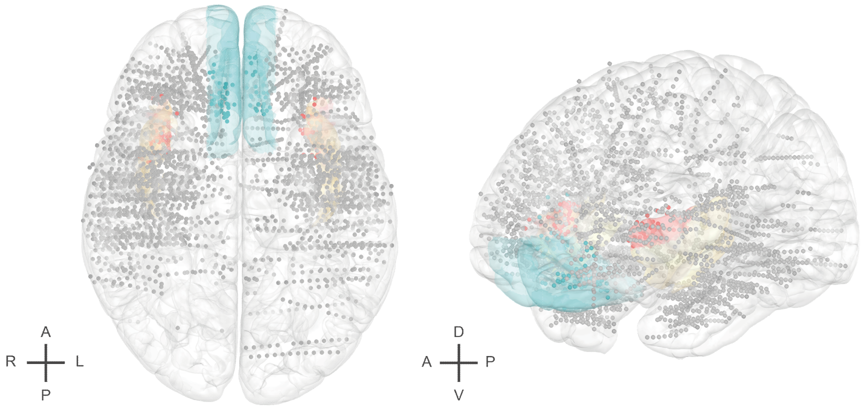


**Figure S1: Anatomical location of all recording sites** (n = 3188 sites acquired from 30 epileptic patients) in the standard Montreal Neurological Institute template brain. Colored brain regions represent the vmPFC (blue) and the daINS (red), and colored dots recording sites located in these region. The whole insula is shown in yellow as reference. Each dot (1 x 1 mm²) represents one recording site (that is, a bipole). Anterior (A), posterior (P), dorsal (D), ventral (V), left (L) and right (R) directions are indicated.

| ID | Sex | Age (years) | Epilepsy onset (age) | Suspected epileptic focus | Hand laterality | Number of electrodes | Number of recording sites | Number of recorded bipoles | Number of bipoles in vmPFC | Number of bipoles in daIns |
|---|---|---|---|---|---|---|---|---|---|---|
| G1 | M | 46 | 23 | Right insulo-opercular / Left opercular | L | 17 | 122 | 85 | 0 | 2 |
| G2 | M | 38 | 15 | Left precentral / Premotor | L | 17 | 122 | 88 | 0 | 1 |
| G3 | M | 43 | 7 | Right temporal | R | 17 | 122 | 83 | 6 | 3 |
| G4 | F | 38 | 3 | Bilateral extensive | R | 18 | 122 | 77 | 0 | 2 |
| G5 | F | 35 | 4 | Several territories | R | 16 | 122 | 92 | 6 | 1 |
| G6 | F | 45 | 10 | Bi-temporal / Amygdala nucleus | R | 17 | 122 | 84 | 6 | 6 |
| G7 | F | 46 | 41 | Right temporal | R | 13 | 122 | 100 | 4 | 2 |
| G8 | F | 43 | 2 | Right insulo-opercular | R | 16 | 122 | 92 | 0 | 3 |
| G9 | M | 45 | 41 | Left mesio-temporal | R | 17 | 122 | 90 | 2 | 4 |
| L1 | F | 33 | 7 | Left insulo-opercular / Left amygdala | R | 13 | 137 | 123 | 2 | 4 |
| L2 | M | 56 | 36 | Left mesio-temporal | R | 13 | 141 | 127 | 2 | 3 |
| L3 | F | 38 | 27 | Right mesio-temporal | R | 9 | 89 | 79 | 2 | 0 |
| M1 | M | 56 | NA | NA | NA | 14 | 183 | 168 | 0 | 3 |
| M2 | M | 34 | NA | NA | NA | 20 | 253 | 228 | 0 | 13 |
| N1 | M | 42 | 6 | Left fronto-opercular | R | 11 | 106 | 93 | 0 | 2 |
| P1 | M | 23 | 15 | Right frontal | R | 14 | 212 | 86 | 9 | 4 |
| P2 | M | 33 | 16 | Left temporal | R | 15 | 126 | 93 | 0 | 1 |
| P3 | F | 46 | 28 | Right temporal | R | 12 | 151 | 136 | 2 | 6 |
| R1 | M | 45 | 10 | Bilateral extensive | R | 13 | 126 | 112 | 3 | 2 |
| R2 | F | 21 | 2 | Right cingulate gyrus | R | 13 | 166 | 152 | 7 | 3 |
| R3 | M | 39 | 30 | Right temporal | R | 13 | 183 | 169 | 6 | 4 |
| R4 | F | 23 | 20 | Right temporo-insulo-perisylvian | A | 11 | 125 | 113 | 0 | 3 |
| R5 | F | 47 | 28 | Hippocampal sclerosis / Right temporal | R | 10 | 107 | 96 | 0 | 2 |
| R6 | M | 17 | 9 | Left temporo-insulo-frontal multifocal | R | 14 | 174 | 159 | 4 | 2 |
| R7 | F | 39 | 8 | Upper posterior frontal gyrus | R | 11 | 126 | 114 | 4 | 2 |
| R8 | M | 46 | 33 | Orbitofrontal / Right anterior temporal | R | 15 | 195 | 179 | 6 | 1 |
| R9 | F | 39 | 26 | Right medial temporal | R | 11 | 124 | 112 | 4 | 2 |
| R10 | M | 21 | 6 | Limbic cingulate | A | 13 | 168 | 154 | 6 | 3 |
| T1 | F | 49 | 33 | Right anterior temporal | R | 13 | 122 | 106 | 5 | 2 |
| T2 | M | 58 | 40 | Left fronto-temporal / Left fronto-mesial | R | 14 | 127 | 104 | 5 | 0 |

**Table S1**: **Demographical data**. M: Male; F: Female; L: Left; R: Right; A: Ambidextrous; vmPFC: Ventromedial Prefrontal Cortex; daIns: Dorsal Anterior Insula; NA: Not available

| | | Mood ratings | | | TML | | | Residual error of choice | | |
|---|---|---|---|---|---|---|---|---|---|---|
| | ROI | Best cluster duration (s) | Sum t-value | p-value | Best cluster duration (s) | Sum t-value | p-value | Best cluster duration (s) | Sum t-value | p-value |
| *Pos.* | **PFCvm** | **0,33** | **122,26** | **0,010** | **0,44** | **132,44** | **$8.10^{-3}$** | **0,33** | **91,20** | **0,020** |
| *Neg.* | Mdm | 0,58 | -153,92 | $7.10^{-3}$ | 4 | -2182,56 | $< 1.7.10^{-5}$ | 0,46 | -114,91 | 0,013 |
| | **daIns** | **0,85** | **-325,84** | **$< 1.7.10^{-5}$** | **0,41** | **-136,38** | **$9.10^{-3}$** | **0,28** | **-85,17** | **0,029** |
| | IPCv | 0,66 | -214,19 | $3.10^{-4}$ | 0,66 | -166,58 | $4.10^{-3}$ | 0,37 | -107,48 | 0,013 |
| | VCrm | 0,37 | -115,37 | 0,022 | 0,28 | -118,65 | 0,025 | 0,44 | -111,13 | 0,016 |

**Table S2: ROIs associated with both mood levels and residual error of choice.** Areas are ordered according to the maximal absolute t-value obtained by averaging best clusters in the three following regression: BGA against mood ratings, BGA against trial-wise mood estimates from the model (TML) and residual error of choice against BGA. Pos: ROI positively associated with mood levels and residual error of choice; Neg: ROIs negatively associated with mood levels and residual error of choice. P-values were obtained with two-sided, one-sample, Student's t-tests cluster-wise corrected ($p_{corr} < 0.05$). vmPFC: Ventromedial Prefrontal Cortex; Mdm: Dorsomedial Motor Cortex; daIns: Dorsal Anterior Insula; IPCv: Ventral Inferior Parietal Cortex; VCrm: Rostral Medial Visual Cortex
